# Extensive hybridization reveals multiple coloration genes underlying a complex plumage phenotype

**DOI:** 10.1101/2020.07.10.197715

**Authors:** Stepfanie M. Aguillon, Jennifer Walsh, Irby J. Lovette

## Abstract

Coloration is an important target of both natural and sexual selection. Discovering the genetic basis of color differences can help us to understand how this visually striking phenotype evolves. Hybridizing taxa with both clear color differences and shallow genomic divergences are unusually tractable for associating coloration phenotypes with their causal genotypes. Here, we leverage the extensive admixture between two common North American woodpeckers—yellow-shafted and red-shafted flickers—to identify the genomic bases of six distinct plumage patches involving both melanin and carotenoid pigments. Comparisons between flickers across ~8.5 million genome-wide SNPs show that these two forms differ at only a small proportion of the genome (mean *F_ST_* = 0.007). Within the few highly differentiated genomic regions, we identify 408 SNPs significantly associated with four of the six plumage patches. These SNPs are linked to multiple genes known to be involved in melanin and carotenoid pigmentation. For example, a gene (CYP2J19) known to cause yellow to red color transitions in other birds is strongly associated with the yellow versus red differences in the wings and tail feathers of these flickers. Additionally, our analyses suggest novel links between known melanin genes and carotenoid coloration. Our finding of patch-specific control of plumage coloration adds to the growing body of literature suggesting color diversity in animals could be created through selection acting on novel combinations of coloration genes.

## INTRODUCTION

Coloration is a visually striking and extraordinarily variable phenotype in animals that drives both natural and sexual selection, and ultimately the process of speciation^1–3^. In recent decades, biologists have been increasingly interested in connecting variation in coloration to an underlying genotype, with much of the focus placed on genes of large effect that influence whole-body coloration differences^3–10^. However, in recent years the use of anonymous genomic scans and admixture mapping has facilitated the discovery of genomic regions involved in coloration of smaller, discrete patches on the body^11–16^. Increasing empirical evidence of patch-specific control of coloration suggests extensive phenotypic diversity could be created through selection acting on novel combinations of coloration genes^17–22^.

Low levels of background genomic divergence—either due to experimental crosses, recent speciation, or ongoing introgression—in taxa that differ primarily in color have allowed for identification of candidate coloration genes in numerous systems^6,14,21,23^. However, what we know about the genes involved in coloration varies extensively depending on the type of pigment involved. The pathways involved in melanin coloration (grays, blacks, browns, and dark reds) are better characterized^5^, compared to carotenoid coloration (bright reds, yellows, and oranges) for which only a handful of underlying genes have been identified^24,25^. This difference is due to differences in pigment acquisition—melanins are produced endogenously, while carotenoids must be acquired through the diet and are subsequently biochemically processed^24^—and the ability to study melanins in humans and other model systems^5^.

Birds with low levels of background divergence have served as particularly powerful non-model systems for discovering the genetic bases of melanin and carotenoid coloration^6,8,11,12,14,18,20,26–28^, as they often exhibit discrete feather patches that differ in color and pigment type across the body^19^. Yet, despite the substantial variation in pigmentation across birds, the genetic bases of melanin and carotenoid coloration have only rarely been studied together in the same system (but see ^11,14,16^), though the genes involved are not currently known to overlap in function or co-localize in the genome^3,5^. Here, we leverage the extensive natural phenotypic variation between yellow-shafted (*Colaptes auratus auratus*) and red-shafted (*C. a. cafer^a^*) flickers, common woodpeckers that hybridize in North America^29^, to identify the genomic underpinnings of plumage coloration and explore the connections between melanin and carotenoid pigmentation. The two flickers differ in the coloration of six distinct feather patches: wings and tail (the eponymous “shaft”), nuchal patch, ear coverts, throat, crown, and male malar stripe (Fig. 1a, Supplementary Table 1)^30^. The pigments involved vary depending on the feather patch, with melanins (throat, ear coverts, crown), carotenoids (wings and tail, nuchal patch), and both melanins and carotenoids (male malar stripe)^31,32^. Previous molecular work has highlighted the very low baseline genetic divergence between these two taxa^33–38^. Importantly, there is extensive ongoing hybridization and backcrossing where the flickers meet in a secondary contact zone in the Great Plains of North America (Supplementary Fig. 1). Admixed and backcrossed hybrids exhibit the full range of possible trait combinations across the six feather patches^30,39,40^.

**Fig. 1.**
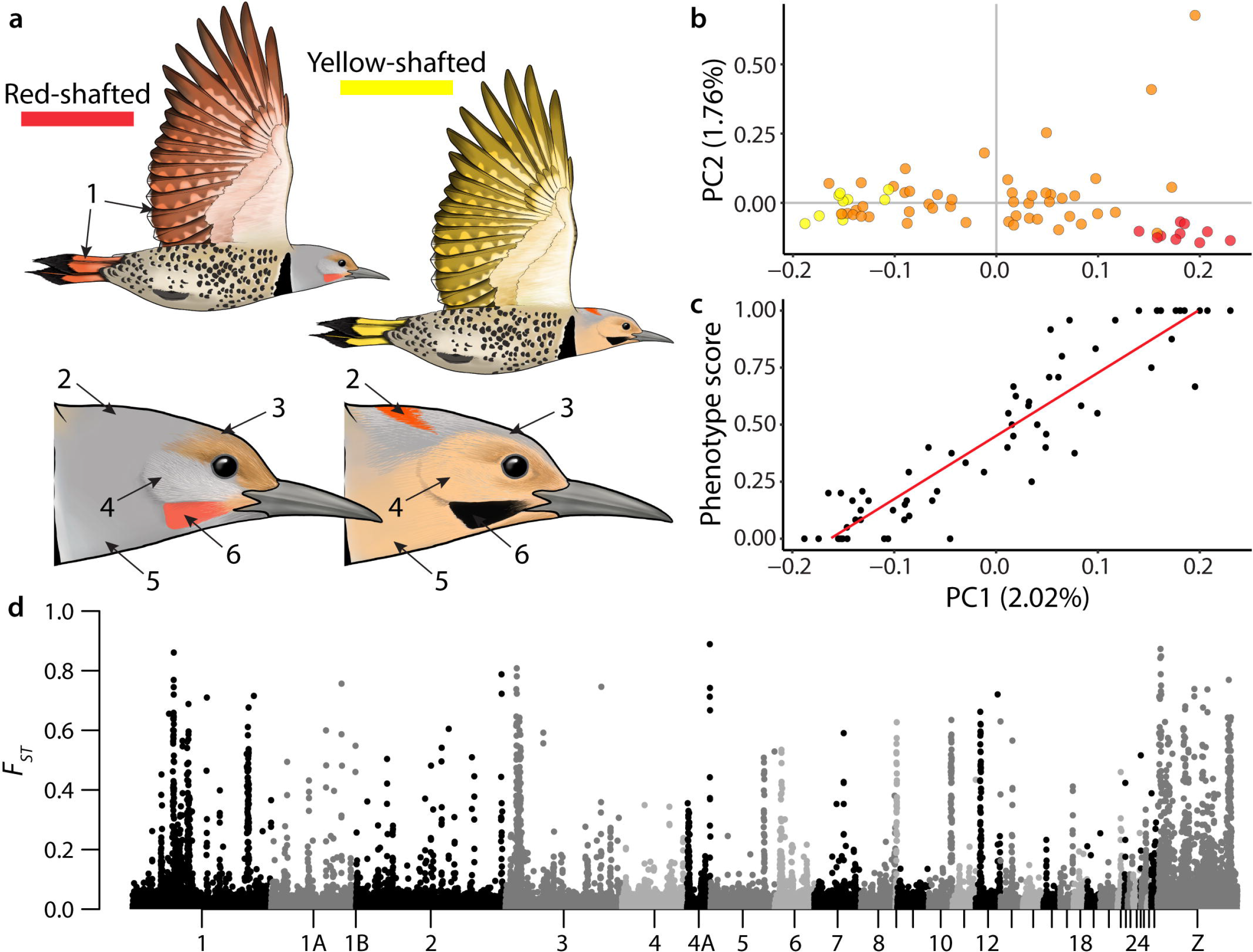
Flicker plumage differences and genomic differentiation. **a**, Coloration differences between red-shafted and yellow-shafted flickers: (1) wings and tail (the eponymous “shaft”), (2) nuchal patch, (3) crown, (4) ear coverts, (5) throat, and (6) male malar stripe. Pigmentation is based on carotenoids (wings and tail, nuchal patch), melanins (crown, ear coverts, throat), and both carotenoids and melanins (male malar stripe). Illustrations by Megan Bishop. **b**, Principal component analysis (PCA) separately clusters yellow-shafted (yellow points), red-shafted (red points), and hybrid (orange points) flickers using the dataset of approximately 8.5 million genome-wide SNPs. **c**, PC1 is significantly associated with overall phenotype score (ρ = 0.93, *P* < 2.2×10^−16^), where variation ranges from 0 for pure yellow-shafted flickers to 1 for pure red-shafted flickers. **d**, The distribution of genetic differentiation (*F_ST_*) between allopatric yellow-shafted flickers and allopatric red-shafted flickers across the whole genome. Individual points show the weighted mean *F_ST_* for 25kb windows. Chromosome positions are based on alignment to the zebra finch genome.

Their combination of low genome-wide divergence, clear phenotypic differences, and extensive hybridization makes flickers an exceptional non-model system in which to explore the genomic basis of feather coloration. Further, the variation in both melanin and carotenoid pigmentation provides an opportunity to explore the potential interactions between genes involved in both pigment types. We compare whole genomes of phenotypically admixed individuals from the hybrid zone along with allopatric red-shafted and yellow-shafted individuals. Here, we (1) assess the genomic landscape of divergence between allopatric flickers and (2) capitalize on a dataset of phenotypically variable hybrid flickers to perform association tests between the genomic markers and the six plumage traits. We leverage these complementary and independent approaches to identify regions of the genome significantly associated with plumage differences. We then (3) search for candidate pigmentation genes present within these genomic regions and (4) discuss potential mechanisms connecting melanin and carotenoid genes with individual plumage patches.

## RESULTS & DISCUSSION

### The genomic landscape of divergence in flickers

We conducted whole genome re-sequencing of 10 allopatric red-shafted, 10 allopatric yellow-shafted, and 48 hybrid flickers (Supplementary Table 2), resulting in approximately 8.5 million SNPs distributed across the genome. Red-shafted and yellow-shafted flickers clustered separately in a principal component analysis (PCA) with hybrids extending between the two parental taxa on PC1 (Fig. 1b; 2.02% of the variation) and clustering separately from them on PC3 and PC4 (Supplementary Fig. 2; 1.69% and 1.67% of the variation, respectively). We estimated *F_ST_* values between the allopatric red-shafted and allopatric yellow-shafted individuals in nonoverlapping 25kb windows to search for divergent regions of the genome. Differentiation across all windows was low between the allopatric individuals (mean genome-wide *F_ST_* = 0.007, mean autosomal *F_ST_* = 0.006, mean Z-linked *F_ST_* = 0.026), but we identified a number of regions with elevated *F_ST_* estimates relative to the background (Fig. 1d). Across the entire dataset, we found only a small number of SNPs that were fixed (790 SNPs with *F_ST_* = 1, 0.009% of the total) or nearly fixed (2,202 SNPs with *F_ST_* > 0.90, 0.026% of the total).

If these highly differentiated regions of the genome contain causal genes related to feather coloration, we expected the PCA to be correlated with phenotypic differences. Thus, we scored the six differing plumage patches (Fig. 1a) in the flickers sampled in the hybrid zone to obtain a score ranging from 0 (yellow-shafted) to 1 (red-shafted). See Methods and Supplementary Table 1 for details on the phenotypic scoring. We found that PC1 was strongly correlated with the overall phenotype score (Fig. 1c, ρ = 0.93, *P* < 2.2×10^−16^). Further, a PCA based on 790 fixed SNPs between allopatric red-shafted and allopatric yellow-shafted flickers resulted in the first PC axis explaining a majority of the variation (55.53%) and individuals spread along PC1 based on overall phenotype score (Supplementary Fig. 3). Taken together, these findings suggest that the few divergent genomic regions between allopatric flickers (*F_ST_* peaks in Fig. 1d) are associated with the loci responsible for their coloration differences.

### Multiple, discrete genomic regions shape the complex plumage phenotype

We took advantage of the plumage trait variation among hybrid flickers to conduct genome-wide associations (GWAs) for each of the six plumage patches to test whether particular *F_ST_* divergence peaks were associated with plumage coloration. By focusing only on hybrid individuals, the results of our GWA analyses are independent from our assessment of genomic divergence between allopatric individuals. Because red-shafted and yellow-shafted flickers are highly ecologically similar^29^ and hybrid flickers with variable trait combinations were sampled from the same geographic transect, we expect any associations identified in the GWAs to be related to differences in plumage coloration traits. 408 SNPs (0.005% of the total) were significantly associated with plumage patches using a significance threshold of α = 0.0000001 (-log_10_(α) = 7), with 25 SNPs identified in more than one analysis (Fig. 2; Supplementary Fig. 4). We found significant associations between multiple SNPs and plumage for four of the six focal traits, excluding throat color (only 1 SNP identified) and crown color (no SNPs identified). We validated our associations to ensure the identified regions represent real associations between plumage patches and genotype using randomized GWA analyses (Supplementary Fig. 5; see Methods for details).

**Fig. 2.**
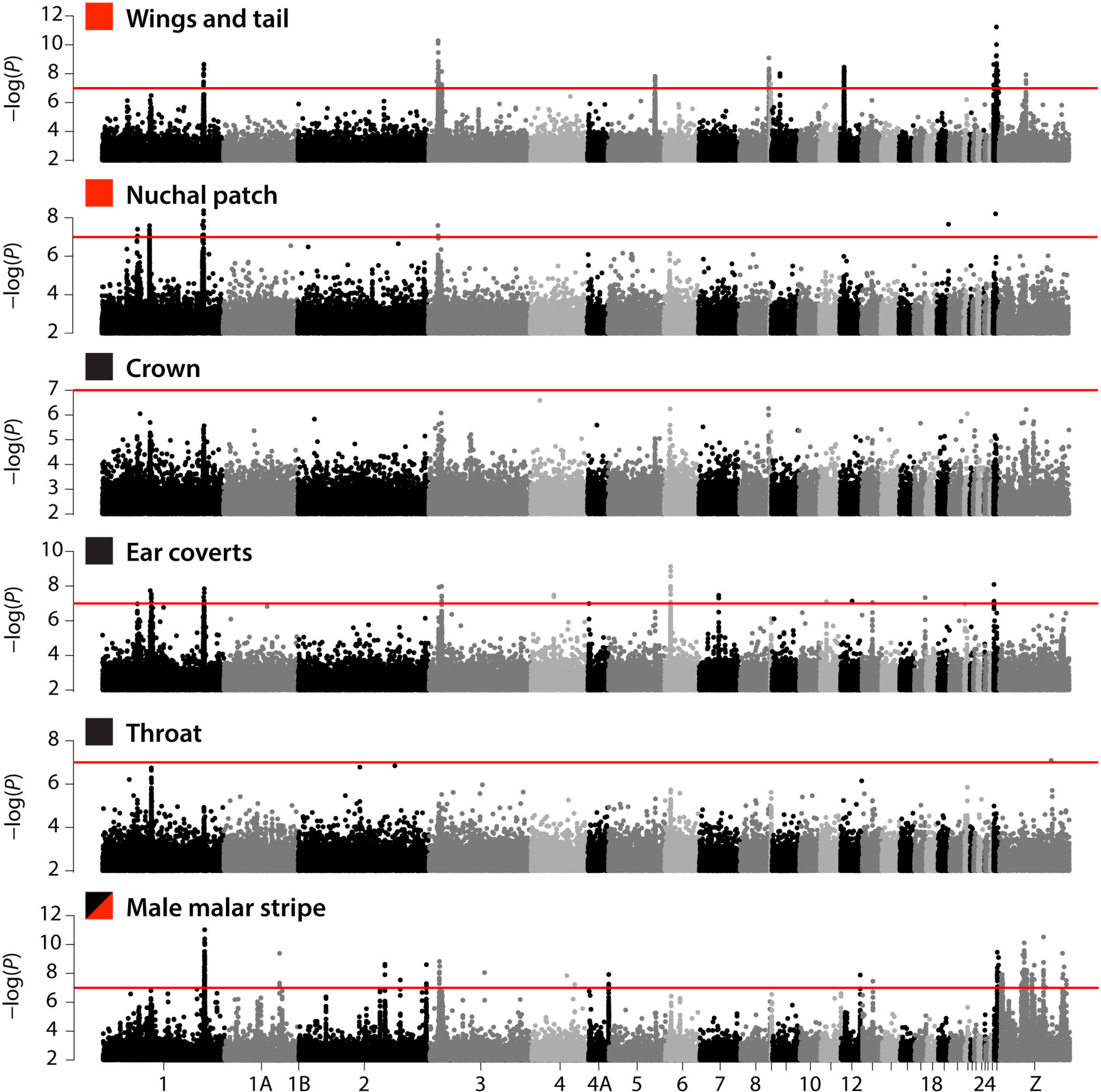
Associations between genomic SNPs and flicker plumage differences. Results from the genome-wide associations (GWAs) of hybrid flickers comparing individual SNPs with coloration differences in the six plumage patches. Pigment type is indicated by the square next to the trait name (red = carotenoid, black = melanin, red and black = carotenoid and melanin). Some peaks of significant SNPs are present in GWAs of multiple phenotypic traits, while other peaks are unique to a single GWA, suggesting multiple mechanisms influence coloration in flickers. The red line represents the significance threshold of −log_10_(*P*) = 7. Chromosome positions are based on alignment to the zebra finch genome. For visualization purposes we show only points with −log_10_(*P*) > 2.

The GWA analyses revealed several genomic regions that were significantly associated with the coloration of the wings and tail, nuchal patch, ear coverts, and male malar stripe (Fig. 2; Supplementary Table 3). In several cases, we identified regions of the genome that were significantly associated with multiple plumage traits (e.g., at the end of chromosome 1 and the beginning of chromosome 3). However, we also identified regions of the genome that were unique to a single GWA analysis (e.g., associations between wings and tail color and regions on chromosomes 5, 8, and 12). These findings suggest multiple mechanisms influencing coloration in flickers: some genomic regions exert pleiotropic control over the coloration of multiple plumage patches, while other genomic regions control the coloration of a single plumage patch (perhaps as loci of large effect). By taking complementary, yet independent, approaches in the GWAs and *F_ST_* analyses, we find that genomic regions identified in the GWAs of hybrid flickers largely lie within regions of the genome with elevated *F_ST_* between allopatric flickers (peaks in Fig. 1d; Supplementary Fig. 6). However, not all genomic regions with elevated *F_ST_* were associated with variation in coloration (e.g., the first peak on chromosome 4A, the peak on chromosome 10, and multiple peaks on the Z chromosome).

### Melanin and carotenoid genes both associate with carotenoid plumage in flickers

To identify candidate genes associated with plumage variation, we searched for all genes within 20kb of genomic regions that were significantly associated with plumage patches (i.e., within the regions identified in Supplementary Table 3). Using this approach, we identified a total of 128 genes (Supplementary Table 4). Here, we highlight 13 genes (Table 1, Fig. 3) that are known or suspected to be involved in melanin or carotenoid pigmentation in other systems: 7 of these candidates are known to be directly involved with pigmentation^6,8,42^, 4 are suspected to be involved in pigmentation based on the function of related genes^8^, and 2 were identified in previous associations with feather coloration in birds^11,26^.

**Table 1.**
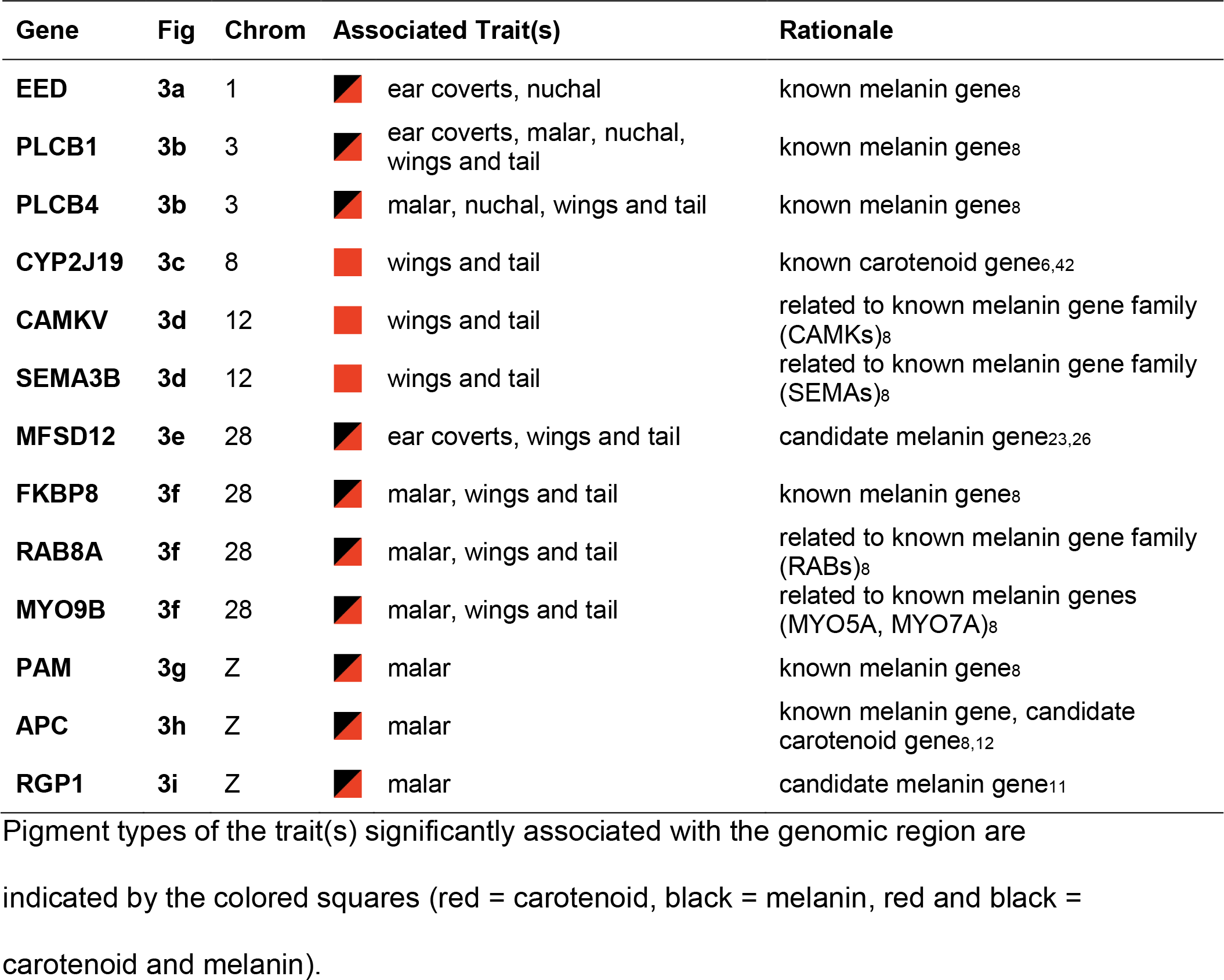
Candidate coloration genes

| Gene | Fig | Chrom | Associated Trait(s) | Rationale |
| --- | --- | --- | --- | --- |
| EED | 3a | 1 | ear coverts, nuchal | known melanin genes<sup>8</sup> |
| PLCB1 | 3b | 3 | ear coverts, malar, nuchal, wings and tail | known melanin genes<sup>8</sup> |
| PLCB4 | 3b | 3 | malar, nuchal, wings and tail | known melanin genes<sup>8</sup> |
| CYP2J19 | 3c | 8 | wings and tail | known carotenoid gene<sup>6,42</sup> |
| CAMKV | 3d | 12 | wings and tail | related to known melanin gene family (CAMKs)<sup>8</sup> |
| SEMA3B | 3d | 12 | wings and tail | related to known melanin gene family (SEMA3s)<sup>8</sup> |
| MFSD12 | 3e | 28 | ear coverts, wings and tail | candidate melanin gene<sup>23,26</sup> |
| FKBP8 | 3f | 28 | malar, wings and tail | known melanin genes<sup>8</sup> |
| RAB8A | 3f | 28 | malar, wings and tail | related to known melanin gene family (RABs)<sup>8</sup> |
| MYO9B | 3f | 28 | malar, wings and tail | related to known melanin genes (MYO5A, MYO7A)<sup>8</sup> |
| PAM | 3g | Z | malar | known melanin genes<sup>8</sup> |
| APC | 3h | Z | malar | known melanin gene, candidate carotenoid gene<sup>8,12</sup> |
| RGP1 | 3i | Z | malar | candidate melanin gene<sup>11</sup> |

indicated by the colored squares (red = carotenoid, black = melanin, red and black = carotenoid and melanin).

**Fig. 3.**
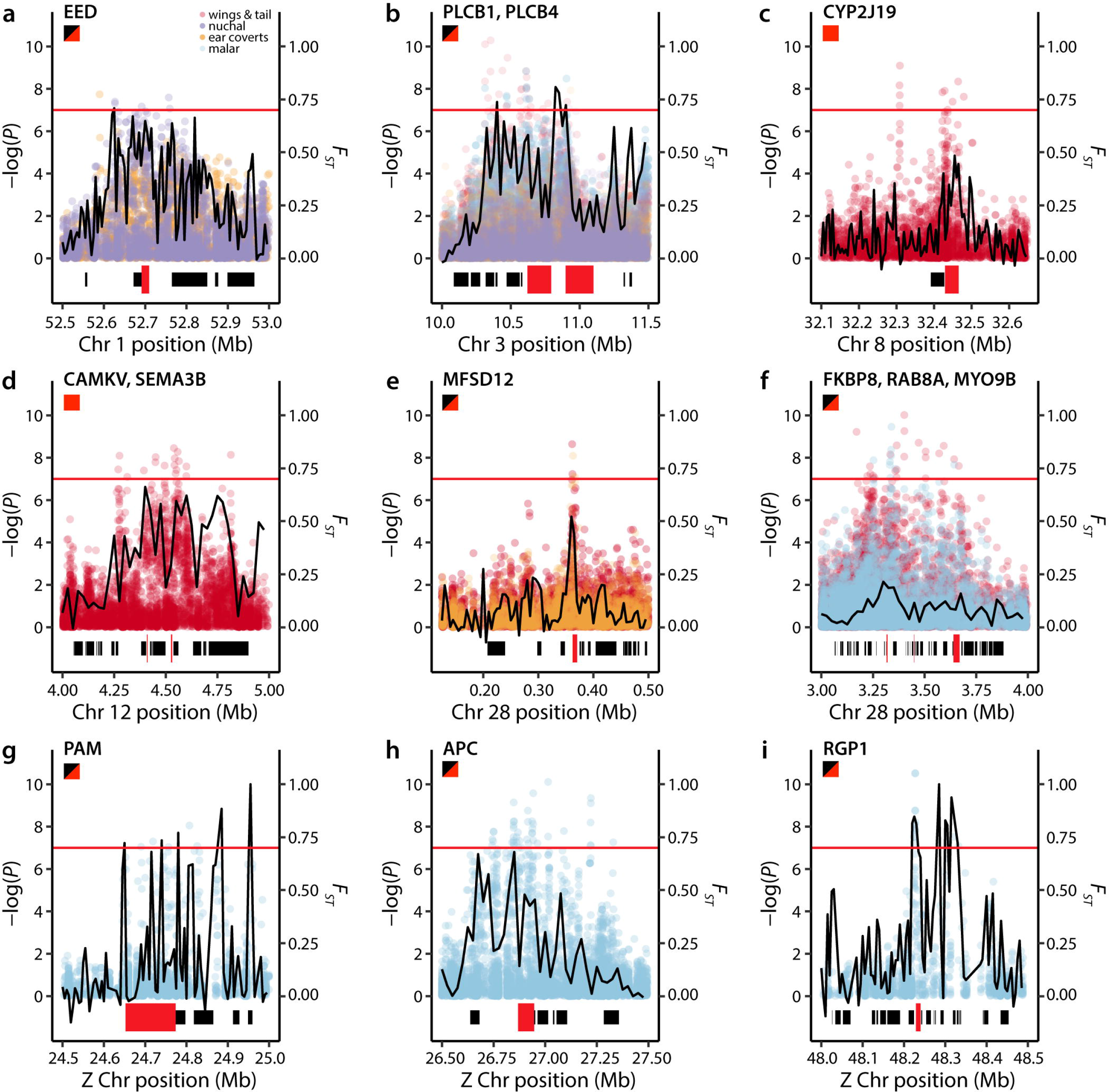
GWA and *F_ST_* patterns around candidate coloration genes. Patterns of genetic differentiation and GWA significance around nine genomic regions of interest containing 13 candidate coloration genes (Table 1). Significance values from the GWA analyses are shown as points colored by the analyses they were identified in (legend in panel **a**). The horizontal red lines indicate the GWA significance threshold of −log_10_(*P*) = 7. Weighted mean *F_ST_* between allopatric red-shafted and allopatric yellow-shafted flickers estimated in 25kb (**b, d, f, h**) or 5kb (**a, c, e, g, i**) windows are shown as black lines in each panel. Genes contained within the plotted area are shown as bars at the bottom of each panel, with the red bars indicating the locations of focal genes. When multiple focal genes are located within a single panel, they are listed at the top of the panel in the order of their physical location (from left to right). Pigment types of the trait(s) significantly associated with the genomic region are indicated by the squares under the gene names (red = carotenoid, black = melanin, red and black = carotenoid and melanin). Chromosome positions are based on alignment to the zebra finch genome.

We find a strong association between wings and tail color and the genomic region on chromosome 8 containing the gene CYP2J19 (Fig. 3c, Table 1), which codes for a cytochrome P450 enzyme. CYP2J19 upregulation via an introgressed variant is causal in changing the typical yellow-feathered canary (*Serinus canaria*) into the “red factor” canary^6^ and the lack of a functional copy in zebra finch (*Taeniopygia guttata*) is implicated in the “yellowbeak” phenotype in which the normally red beak and legs are instead yellow^42^. It is currently one of only two genes known to be involved in red coloration in birds, but evidence of its functioning in natural systems is still limited (but see ^43–45^). Our identification of CYP2J19 in the GWA for wings and tail coloration suggests that it mediates this yellow versus red trait difference in flickers and provides further support for its importance in red coloration across diverse avian lineages.

The majority of our identified candidate genes for carotenoid plumage patches in the flickers (Table 1) are known or suspected to affect melanin pigmentation in other organisms (henceforth “melanin genes”). In some cases, we find melanin genes are associated with both melanin and carotenoid traits in a single region of the genome (EED on chromosome 1 (Fig. 3a), PLCB1 and PLCB4 on chromosome 3 (Fig. 3b), and MFSD12 (Fig. 3e) and FKBP8, RAM8A, and MYO9B (Fig. 3f) on chromosome 28), while in other cases, we find melanin genes associated with a single trait (malar stripe) that uses both pigment types (PAM (Fig. 3g), APC (Fig. 3h), and RGP1 (Fig. 3i) on the Z chromosome). Most unusually, we identify a region on chromosome 12 associated only with differences in wings and tail color—a carotenoid-based trait—that contains CAMKV and SEMA3B (Fig. 3d), which are both related to known melanin pathway gene families. To our knowledge, of these 12 melanin genes only APC has previously been linked to carotenoid pigmentation (in an associational study)^12^, in addition to its known link to melanin pigmentation^8^.

### Potential mechanisms linking melanin genes with carotenoid traits

Melanin and carotenoid pigmentation derive from different biochemical pathways^46^ and the genes involved in the different processes are not currently known to co-localize in the genome or exert influence over each other^3,5^. Thus, our finding in flickers of repeated associations between different carotenoid traits and melanin genes was unexpected. Although we lack a complete annotation of the flicker reference genome and therefore may have missed some causal genes, we repeatedly found associations between known carotenoid traits and melanin genes from different regions of the genome. Here, we outline three non-mutually exclusive explanations for these associations that link melanin genes with carotenoid plumage.

First, melanin genes could be associated with carotenoid traits because the trait differences we observe are actually due to a combination of both pigments. Melanin genes associated with the male malar stripe (Fig. 3b, Fig. 3f-i) exemplify this mechanism: red pigments are present in the malar stripes of both red-shafted and yellow-shafted flickers, and yellow-shafted flickers subsequently overlay melanin to produce a black malar stripe that masks the red pigment^31,47^. Beyond this one situation where melanic pigments completely overlay a carotenoid trait, it is also possible that the two pigments are used in concert within the feathers to produce the observed color (as in ^48,49^). Additionally, melanins serve a number of other functions in feathers apart from coloration (e.g., feather structure and stability^50,51^, UV protection^52^, resistance to bacterial degradation^53^), so differences in the feathers beyond coloration could also exist.

Second, some of the association patterns we identify in the GWAs suggest pleiotropic effects of melanin genes: we find multiple plumage patches (carotenoid and melanin) associated with the same region of the genome (Fig. 3a-b, Fig. 3e-f). Because we find different combinations of traits associated with different genomic regions—rather than overlapping patterns across all analyses—the results support some genomic regions exerting control over multiple plumage patches. This could occur through regulatory genes typically involved in melanin pigmentation evolving to control the expression of both melanins and carotenoids. Similar results have been found in two different warbler species (*Setophaga*), where associations between a single genomic region and multiple aspects of carotenoid and melanin plumage differences have been identifed^11,16^. The finding of pleiotropic effects on melanin and carotenoid plumage in woodpeckers and warblers, distantly related bird taxa, suggests pleiotropic effects of melanin genes may be widespread.

Finally, our results suggest the intriguing possibility that genes typically thought to be involved in melanin pigmentation could also be used in creating carotenoid traits. In particular, the association between wings and tail color with CAMKV and SEMA3B (Fig. 3d), suggests the potential for melanin genes to play a role in carotenoid traits. Rather than controlling the upregulation and deposition of melanin (as in our first explanation), these genes may instead control the absence of melanin within the feathers, as a reduction of melanin is necessary for the bright red and yellow coloration to be visible. It is possible that the two taxa have differential levels of melanin reduction or that they use different molecular pathways. This finding opens up a novel area of inquiry aimed at understanding the interactions between melanin genes and the production and display of carotenoid traits. Exploring differences in gene expression in these colored feather patches could help to better understand the mechanisms underlying these associations.

### Conclusions

Here, we identify a complex relationship linking melanin and carotenoid genes with modular plumage patches. We provide evidence for a novel link between known melanin genes and carotenoid traits. Additionally, we identified CYP2J19 as a strong candidate related to red versus yellow coloration differences, providing further evidence of its action across divergent bird lineages. The extensive hybridization between red-shafted and yellow-shafted flickers, in combination with their clear phenotypic differences, has allowed us to separately connect phenotypic differences with individual genomic regions. The patch-specific control of plumage coloration that we identify here, and increasingly found in other systems^19^, suggests color diversity across birds could be created through selection to produce novel combinations of coloration genes each exerting control on a separate body patch.

## METHODS

### Sample collection and plumage scoring

We obtained tissue samples from allopatric yellow-shafted flickers from New York (*N* = 5) and Florida (*N* = 5), and allopatric red-shafted flickers from Oregon (*N* = 5) and California (*N* = 5). These allopatric samples allowed us to characterize genomic differentiation between the flickers far from the region of current hybridization. Additionally, we sampled flickers with variable phenotypes (*N* = 48) from a sampling transect across the hybrid zone in Nebraska and Colorado following the Platte River. See Supplementary Table 2 for details on included samples.

Red-shafted and yellow-shafted flickers differ in color across six distinct plumage characters: wings and tail, nuchal patch, crown, ear coverts, throat, and male malar stripe (Fig. 1a)^29^. Hybrids exhibit various combinations of parental traits and traits intermediate to the parental traits. We scored plumage characters on a scale from 0 (pure yellow-shafted) to 4 (pure red-shafted) following a protocol slightly modified from Short^30^ (Supplementary Table 1). We additionally calculated an overall phenotype score by adding the scores for the six individual traits and standardizing to range from 0 to 1 to include both sexes (as all females lack a malar stripe). To ensure consistency, all scoring was conducted by a single individual (SMA). Hybrid flickers were chosen for genotyping in this study to maximize power in the GWA analyses: we selected a panel of hybrids that exhibited high variation in their combination of plumage traits.

### Reference genome assembly and annotation

We sequenced and assembled the genome of a male yellow-shafted flicker (CUMV 57446). DNA was extracted using the Gentra Puregene Tissue Kit following the manufacturer’s protocol (Qiagen, California, USA) to isolate high molecular weight DNA. Three libraries were prepared and sequenced by the Cornell Weill Medical College genomics core facility—one 180bp fragment library and two mate-pair libraries (3kb and 8kb insert size)—across three lanes of an Illumina HiSeq2500 using the rapid run mode. The two mate-pair libraries were multiplexed on a single lane, while the fragment library was run across two lanes. The three lanes of sequencing generated ~481 million raw paired-end reads.

We assembled the reference genome using ALLPATHS-LG v.52488^54^ and assessed the quality of the assembly using QUAST v.4.0^55^ and BUSCO v.3.1.0^56^. The reference assembly had a total length of 1.10 Gb distributed across 22,654 scaffolds with an N50 of 1.57 Mb. Using BUSCO, we searched for a set of 2,586 conserved, single-copy orthologs found across vertebrates. Our flicker reference genome contained a single, complete copy of 87.2% of these genes and a fragment of an additional 6.7%. Of the remaining genes, 0.4% were identified multiple times and 5.7% were completely missing. To obtain more precise information on chromosome position, we additionally assigned individual scaffolds to chromosomes based on assignments in the Ensembl zebra finch (*Taeniopygia guttata*) reference genome v.3.2.4 release 91^57^ using the ‘Chromosemble’ function in Satsuma^58^.

We annotated the reference genome using the MAKER v.2.31 pipeline^59^. We first used RepeatModeler v.1.0^60^ to generate a library of repetitive sequences present in the assembly and RepeatMasker v.4.0^61^ to soft mask these repeats. We then produced gene models by running two iterations of MAKER: the first iteration produces ab initio gene predictions, while the second iteration uses the gene models predicted from the first to improve performance. We used the Ensembl expressed sequence tags (ESTs) and protein database from the zebra finch (v.3.2.4 downloaded July 2017)^57^ to train MAKER. This pipeline annotated a total of 12,141 genes (62.4% of the total). 97.3% of the proteins predicted by MAKER matched zebra finch proteins using BLAST^62^.

### Low coverage re-sequencing and variant discovery

We performed low coverage re-sequencing of the genomes of 68 flickers. Genomic DNA was extracted from each sample using DNeasy blood and tissue extraction kits (Qiagen, California, USA) and DNA concentrations were determined using a Qubit fluorometer (Life Technologies, California, USA). We used 200 ng of DNA from each sample to prepare individually barcoded libraries with a 550bp insert size following the protocol for the TruSeq Nano DNA Library Prep kit (Illumina, California, USA). The libraries were pooled into three groups and sequenced separately on an Illumina NextSeq500 lane (2×150bp) at the Cornell University Biotechnology Resource Center.

We assessed the quality of individual libraries using FastQC v.0.11.5 (http://www.bioinformatics.babraham.ac.uk/projects/fastqc/) and subsequently performed trimming, adapter removal, and initial quality filtering using AdapterRemoval v.2.1.^63^. We required a minimum Phred quality score of 10 and merged overlapping paired-end reads. Filtered sequences were aligned to the northern flicker reference genome using Bowtie2 v.2.2.8^64^ with the very sensitive, local option. Alignment statistics were obtained from Qualimap v.2.2.1^65^; the average alignment rate across all samples was 92.3%. Alignment rates were comparable across the different taxa: red-shafted flickers (91.9%), yellow-shafted flickers (91.8%), and hybrid flickers (92.6%). After filtering and aligning to the reference, the mean depth of coverage was 5.8X (range: 2.0X – 15.1X).

All resulting SAM files were converted to BAM files, sorted, and indexed using SAMtools v.1.3^66^. We used Picard Tools v.2.8.2 (https://broadinstitute.github.io/picard/) to mark PCR duplicates and subsequently realigned around indels and fixed mate-pairs using GATK v.3.8.1^67^. Variant discovery and genotyping for the 68 flickers was performed using the unified genotyper module in GATK. We used the following hard filtering parameters to remove variants from the output file: QD < 2.0, FS > 40.0, MQ < 20.0, and HaplotypeScore > 12.0. Subsequently, we filtered out variants that were not biallelic, had a minor allele frequency less than 5%, had a mean depth of coverage less than 3X or greater than 50X, and had more than 20% missing data across all individuals in the dataset. This pipeline produced 8,495,326 SNPs genotyped across all 68 flickers. We repeated the analyses with a variety of other SNP calling tools, including ANGSD^68^ and the haplotype caller module in GATK^67^. We obtained qualitatively similar results across all analyses, and so here choose to present results from SNP calling with unified genotyper in GATK.

### Population genomic analyses

We visualized genetic clustering in the SNP dataset by performing a principal component analysis (PCA) using the ‘snpgdsPCA’ function in the SNPRelate package^69^ in R v.3.5.2^70^. We characterized genome-wide patterns of divergence between allopatric red-shafted and allopatric yellow-shafted flickers by calculating *F_ST_* using VCFtools v.0.1.16^71^ across 5 and 25 kb windows and individual SNPs. We visualized the results using the ‘manhattan’ function in the qqman package^72^ in R. Additionally, we used VCFtools to calculate nucleotide diversity, heterozygosity, and Tajima’s D (Supplementary Table 5).

### Genotype-phenotype associations

We used GEMMA v.0.98 (Genome-wide Efficient Mixed Model Association)^73^ to associate genotypic variation at SNPs with variation in the six plumage traits for the 48 hybrid flickers while controlling for levels of relatedness. The GEMMA analysis requires a complete SNP dataset, so we first used BEAGLE v.4.1^74^ to impute missing data in the dataset. We transformed the imputed dataset into binary PLINK BED format using VCFtools v.0.1.16^71^ and PLINK v. 1.09^75^. We calculated a relatedness matrix in GEMMA using the centered relatedness matrix option (-gk 1). We conducted separate univariate linear mixed models for each phenotypic trait and used the Wald test (-lmm 1) with a significance threshold of α = 0.0000001 (-log_10_(α) = 7) to identify significant associations between SNPs and phenotypes. To visualize the results, we used the ‘manhattan’ function in the qqman package^72^ in R.

To validate the resulting associations, we also repeated the GEMMA analysis using a dataset with randomized phenotypes. Instead of generating artificial phenotypic scores, we retained the true phenotypic scoring across all plumage traits, but randomized the individual assignment. If the GEMMA analysis was identifying real associations between genotype and phenotype, we expected few SNPs to exceed our significance threshold in this randomized analysis. In strong contrast to the non-randomized results, we found only six significant SNPs across the six randomized analyses and no clustering of significant SNPs in any genomic region (Supplementary Fig. 5).

### Functional characterization of candidate genes

We compiled a list of genes within a 20 kb buffer of SNPs significantly associated with phenotype using Geneious v.11.1.5^76^. To characterize putative candidate genes, we used ontology information from the zebra finch Ensembl database^57^ and functional information from the Uniprot database^77^. We additionally compared the identified list of genes to known genes involved in pigmentation. We were able to compare our gene list to 428 genes known to be involved in melanin pigmentation^8^, and searched for the three gene families known to be involved in carotenoid pigmentation (β-carotene oxygenases, scavenger receptors, and cytochrome P450s)^24^ and genes identified in recent analyses of pigmentation in other bird species^11,12,14,16,26^.

## ACKNOWLEDGMENTS

SMA would like to acknowledge the pivotal role the late Richard G. Harrison played in developing her dissertation project. The authors thank the Cornell University Museum of Vertebrates, Burke Museum, and Louisiana State Museum of Natural Science for contributing samples for this work, as well as the many collectors who contribute samples to museums. We thank V. G. Rohwer for collecting many of the samples used in this study and for coordinating field work logistics. B. Mims, N. A. Kramer, and T. Brooks assisted with field work. We thank B. G. Butcher for guidance and assistance with laboratory work, L. Campagna for assistance with reference genome sequencing and assembly, and D. P. L. Toews for input on bioinformatics analyses. We appreciate the insightful comments provided by members of the Fuller Evolutionary Biology Program, L. Campagna, G. Hill, J. Hudon, N. Justyn, M. Powers, and D. P. L. Toews on earlier versions of this manuscript. We thank Megan Bishop for providing illustrations of the flickers.

## DATA AVAILABILITY

Reference genome and re-sequencing data will be made available upon manuscript acceptance. Scripts for analyses are available at: https://github.com/stepfanie-aguillon/flicker-WGS.

## AUTHOR CONTRIBUTIONS

SMA and IJL conceived the study. SMA collected the data. SMA analyzed the data with input from JW. SMA, JW, and IJL wrote and reviewed the manuscript.

## Footnotes

a Note to reviewers and Editor: The subspecific epithet of the red-shafted flicker is etymologically based on a term referring to an African people that is an extreme racial slur. This nomenclatural history places users of this official Linnaean name in the unfortunate situation of perpetuating this slur. We include the official Linnaean name in this one line, but otherwise purposefully refer to these taxa by their common names. Aguillon and Lovette have elsewhere proposed the scientific name for the red-shafted flicker be changed to *Colaptes auratus lathami*, but this name is not yet widely accepted^41^.

## REFERENCES

1 Cuthill, I. C. et al. The biology of color. Science 357, eaan0221 (2017).

2 Hill, G. E. & McGraw, K. J. Bird Coloration: Function and Evolution. Vol. 2 (Harvard University Press, 2006).

3 Orteu, A. & Jiggins, C. D. The genomics of coloration provides insights into adaptive evolution. Nat. Rev. Genet. (2020).

4 Hoekstra, H. E., Hirschmann, R. J., Bundey, R. A., Insel, P. A. & Crossland, J. P. A single amino acid mutation contributes to adaptive beach mouse color pattern. Science 313, 101–104 (2006).

5 Hubbard, J. K., Uy, J. A., Hauber, M. E., Hoekstra, H. E. & Safran, R. J. Vertebrate pigmentation: from underlying genes to adaptive function. Trends Genet. 26, 231–239 (2010).

6 Lopes, R. J. et al. Genetic basis for red coloration in birds. Curr. Biol. 26, 1427–1434 (2016).

7 Mundy, N. I. A window on the genetics of evolution: MC1R and plumage colouration in birds. Proc. Biol. Sci. 272, 1633–1640 (2005).

8 Poelstra, J. W., Vijay, N., Hoeppner, M. P. & Wolf, J. B. Transcriptomics of colour patterning and coloration shifts in crows. Mol. Ecol. 24, 4617–4628 (2015).

9 San-Jose, L. M. & Roulin, A. Genomics of coloration in natural animal populations. Phil. Trans. R. Soc. B Biol. Sci. 372, 20160337 (2017).

10 Uy, J. A. et al. Mutations in different pigmentation genes are associated with parallel melanism in island flycatchers. Proc. Biol. Sci. 283, 20160731 (2016).

11 Brelsford, A., Toews, D. P. L. & Irwin, D. E. Admixture mapping in a hybrid zone reveals loci associated with avian feather coloration. Proc. Biol. Sci. 284, 20171106 (2017).

12 Gao, G. et al. Comparative genomics and transcriptomics of *Chrysolophus* provide insights into the evolution of complex plumage coloration. Gigascience 7, giy113 (2018).

13 Kim, K. W. et al. Genetics and evidence for balancing selection of a sex-linked colour polymorphism in a songbird. Nat. Commun. 10, 1852 (2019).

14 Toews, D. P. et al. Plumage genes and little else distinguish the genomes of hybridizing warblers. Curr. Biol. 26, 2313–2318 (2016).

15 Toomey, M. B. et al. A non-coding region near Follistatin controls head colour polymorphism in the Gouldian finch. Proc. Biol. Sci. 285, 20181788 (2018).

16 Wang, S. et al. Selection on a pleiotropic color gene block underpins early differentiation between two warbler species. Preprint at https://doi.org/10.1101/853390 (2019).

17 Mazo-Vargas, A. et al. Macroevolutionary shifts of WntA function potentiate butterfly wing-pattern diversity. Proc. Natl. Acad. Sci. USA 114, 10701–10706 (2017).

18 Campagna, L. et al. Repeated divergent selection on pigmentation genes in a rapid finch radiation. Sci. Adv. 3, e1602404 (2017).

19 Funk, E. R. & Taylor, S. A. High-throughput sequencing is revealing genetic associations with avian plumage color. Auk 136, 1–7 (2019).

20 Stryjewski, K. F. & Sorenson, M. D. Mosaic genome evolution in a recent and rapid avian radiation. Nat. Ecol. Evol. 1, 1912–1922 (2017).

21 Van Belleghem, S. M. et al. Complex modular architecture around a simple toolkit of wing pattern genes. Nat. Ecol. Evol. 1, 0052 (2017).

22 Zhang, L., Mazo-Vargas, A. & Reed, R. D. Single master regulatory gene coordinates the evolution and development of butterfly color and iridescence. Proc. Natl. Acad. Sci. USA 114, 10707–10712 (2017).

23 Rodriguez, A., Mundy, N. I., Ibanez, R. & Prohl, H. Being red, blue and green: the genetic basis of coloration differences in the strawberry poison frog (*Oophaga pumilio*). BMC Genomics 21, 301 (2020).

24 Toews, D. P., Hofmeister, N. R. & Taylor, S. A. The Evolution and Genetics of Carotenoid Processing in Animals. Trends Genet. 33, 171–182 (2017).

25 Walsh, N., Dale, J., McGraw, K. J., Pointer, M. A. & Mundy, N. I. Candidate genes for carotenoid coloration in vertebrates and their expression profiles in the carotenoid-containing plumage and bill of a wild bird. Proc. Biol. Sci. 279, 58–66 (2012).

26 Abolins-Abols, M. et al. Differential gene regulation underlies variation in melanic plumage coloration in the dark-eyed junco (*Junco hyemalis*). Mol. Ecol. 27, 4501–4515 (2018).

27 Cooke, T. F. et al. Genetic Mapping and Biochemical Basis of Yellow Feather Pigmentation in Budgerigars. Cell 171, 427–439 e421 (2017).

28 Gazda, M. A. et al. A genetic mechanism for sexual dichromatism in birds. Science 368, 1270–1274 (2020).

29 Wiebe, K. L. & Moore, W. S. Northern Flicker (*Colaptes auratus*) in The Birds of North America (Cornell Lab of Ornithology, 2017).

30 Short, L. L. Hybridization in the flickers (*Colaptes*) of North America. Bull. Am. Mus. Nat. Hist. N. Y. 129, 307–428 (1965).

31 Hudon, J., Wiebe, K. L., Pini, E. & Stradi, R. Plumage pigment differences underlying the yellow-red differentiation in the Northern Flicker (*Colaptes auratus*). Comp. Biochem. Phys. B 183C, 1–10 (2015).

32 Wiebe, K. L. & Vitousek, M. N. Melanin plumage ornaments in both sexes of Northern Flicker are associated with body condition and predict reproductive output independent of age. Auk 132, 507–517 (2015).

33 Aguillon, S. M., Campagna, L., Harrison, R. G. & Lovette, I. J. A flicker of hope: Genomic data distinguish Northern Flicker taxa despite low levels of divergence. Auk 135, 748–766 (2018).

34 Fletcher, S. D. & Moore, W. S. Further analysis of allozyme variation in the Northern Flicker, in comparison with mitochondrial DNA variation. Condor 94, 988–991 (1992).

35 Grudzien, T. A. & Moore, W. S. Genetic differentiation between the yellow-shafted and red-shafted subspecies of the northern flicker. Biochem. Syst. Ecol. 14, 451–453 (1986).

36 Grudzien, T. A., Moore, W. S., Cook, J. R. & Tagle, D. Genic population structure and gene flow in the northern flicker (*Colaptes auratus*) hybrid zone. Auk 104, 654–664 (1987).

37 Manthey, J. D., Geiger, M. & Moyle, R. G. Relationships of morphological groups in the northern flicker superspecies complex (*Colaptes auratus* & *C. chrysoides*). Syst. Biodivers. 15, 183–191 (2017).

38 Moore, W. S., Graham, J. H. & Price, J. T. Mitochondrial DNA variation in the northern flicker (*Colaptes auratus*, Aves). Mol. Biol. Evol. 8, 327–344 (1991).

39 Moore, W. S. & Buchanan, D. B. Stability of the northern flicker hybrid zone in historical times: implications for adaptive speciation theory. Evolution 39, 135–151 (1985).

40 Moore, W. S. Random mating in the northern flicker hybrid zone: implications for the evolution of bright and contrasting plumage patterns in birds. Evolution 41, 539–546 (1987).

41 Aguillon, S. M. & Lovette, I. J. Change the specific/subspecific/morphological group name of the Red-shafted Flicker from *cafer* to *lathami*. AOS North American Classification Committee 2019-A-10, 66–71 (2019).

42 Mundy, N. I. et al. Red carotenoid coloration in the zebra finch is controlled by a Cytochrome P450 gene cluster. Curr. Biol. 26, 1435–1440 (2016).

43 Hooper, D. M., Griffith, S. C. & Price, T. D. Sex chromosome inversions enforce reproductive isolation across an avian hybrid zone. Mol. Ecol. 28, 1246–1262 (2019).

44 Kirschel, A. et al. CYP2J19 mediates carotenoid colour introgression across a natural avian hybrid zone. Preprint at https://doi.org/10.22541/au.158880215.54508683/v2 (2020).

45 Twyman, H., Prager, M., Mundy, N. I. & Andersson, S. Expression of a carotenoid-modifying gene and evolution of red coloration in weaverbirds (Ploceidae). Mol. Ecol. 27, 449–458 (2018).

46 Hill, G. E. & McGraw, K. J. Bird Coloration: Mechanisms and Measurements. Vol. 1 (Harvard University Press, 2006).

47 Test, F. H. The nature of red, yellow, and orange pigments in woodpeckers of the genus *Colaptes*. Univ. Calif. Publ. Zool. 46, 371–390 (1942).

48 Hofmann, C. M., McGraw, K. J., Cronin, T. W. & Omland, K. E. Melanin coloration in New World orioles I: carotenoid masking and pigment dichromatism in the orchard oriole complex. J. Avian Biol. 38, 163–171 (2007).

49 McGraw, K. J., Wakamatsu, K., Clark, A. B. & Yasukawa, K. Red-winged blackbirds *Agelaius phoeniceus* use carotenoids and melanin pigments to color their epaulets. J. Avian Biol. 35, 543–550 (2004).

50 Bonser, R. H. C. Melanin and the abrasion resistance of feathers. Condor 97, 590–591 (1995).

51 Kose, M. & Møller, A. P. Sexual selection, feather breakage and parasites: the importance of white spots in the tail of the barn swallow (*Hirundo rustica*). Behav. Ecol. Sociobiol. 45, 430–436 (1999).

52 Ward, J. M., Blount, J. D., Ruxton, G. D. & Houston, D. C. The adaptive significance of dark plumage for birds in desert environments. Ardea 90, 311–323 (2002).

53 Peele, A. M., Burtt, E. H., Jr., Schroeder, M. R. & Greenberg, R. S. Dark color of the coastal plain swamp sparrow (*Melospiza georgiana nigrescens*) may be an evolutionary response to occurrence and abundance of salt-tolerant feather-degrading bacillin in its plumage. Auk 126, 531–535 (2009).

54 Gnerre, S. et al. High-quality draft assemblies of mammalian genomes from massively parallel sequence data. Proc. Natl. Acad. Sci. USA 108, 1513–1518 (2011).

55 Gurevich, A., Saveliev, V., Vyahhi, N. & Tesler, G. QUAST: quality assessment tool for genome assemblies. Bioinformatics 29, 1072–1075 (2013).

56 Simão, F. A., Waterhouse, R. M., Ioannidis, P., Kriventseva, E. V. & Zdobnov, E. M. BUSCO: assessing genome assembly and annotation completeness with single-copy orthologs. Bioinformatics 31, 3210–3212 (2015).

57 Aken, B. L. et al. Ensembl 2017. Nucleic Acids Res. 45, D635–D642 (2017).

58 Grabherr, M. G. et al. Genome-wide synteny through highly sensitive sequence alignment: Satsuma. Bioinformatics 26, 1145–1151 (2010).

59 Cantarel, B. L. et al. MAKER: an easy-to-use annotation pipeline designed for emerging model organism genomes. Genome Res. 18, 188–196 (2008).

60 RepeatModeler Open-1.0 (2008).

61 RepeatMasker Open-4.0 (2013).

62 Altschul, S. F., Gish, W., Miller, W., Myers, E. W. & Lipman, D. J. Basic Local Alignment Search Tool. J. Mol. Biol. 215, 403–410 (1990).

63 Lindgreen, S. AdapterRemoval: easy cleaning of next-generation sequencing reads. BMC Res. Notes 5, 337 (2012).

64 Langmead, B. & Salzberg, S. L. Fast gapped-read alignment with Bowtie 2. Nat. Methods 9, 357–359 (2012).

65 Okonechnikov, K., Conesa, A. & Gárcia-Alcalde, F. Qualimap 2: advanced multisample quality control for high-throughput sequencing data. Bioinformatics 32, 292–294 (2016).

66 Li, H. et al. The Sequence Alignment/Map format and SAMtools. Bioinformatics 25, 2078–2079 (2009).

67 McKenna, A. et al. The Genome Analysis Toolkit: a MapReduce framework for analyzing next-generation DNA sequencing data. Genome Res. 20, 1297–1303 (2010).

68 Korneliussen, T. S., Albrechtsen, A. & Nielsen, R. ANGSD: Analysis of Next Generation Sequencing Data. BMC Bioinformatics 15, 356 (2014).

69 Zheng, X. et al. A high-performance computing toolset for relatedness and principal component analysis of SNP data. Bioinformatics 28, 3326–3328 (2012).

70 R: A language and environment for statistical computing (R Foundation for Statistical Computing, Vienna, Austria, 2018).

71 Danecek, P. et al. The variant call format and VCFtools. Bioinformatics 27, 2156–2158 (2011).

72 Turner, S. D. qqman: an R package for visualizing GWAS results using Q-Q and manhattan plots. Preprint at https://doi.org/10.1101/005165 (2014).

73 Zhou, X. & Stephens, M. Genome-wide efficient mixed-model analysis for association studies. Nat. Genet. 44, 821–824 (2012).

74 Browning, S. R. & Browning, B. L. Rapid and accurate haplotype phasing and missing-data inference for whole-genome association studies by use of localized haplotype clustering. Am. J. Hum. Genet. 81, 1084–1097 (2007).

75 Purcell, S. et al. PLINK: a toolset for whole-genome association and populationbased linkage analysis. Am. J. Hum. Genet. 81, 559–575 (2007).

76 Kearse, M. et al. Geneious Basic: an integrated and extendable desktop software platform for the organization and analysis of sequence data. Bioinformatics 28, 1647–1649 (2012).

77. UniProt Consortium. UniProt: a worldwide hub of protein knowledge. Nucleic Acids Res. 47, D506–D515 (2019).

