## Supplemental Information for "Extensive hybridization reveals multiple coloration genes underlying a complex plumage phenotype"

**Supplementary Table 1 | Flicker plumage traits and phenotypic scoring of hybrids.** Details on the six plumage patch coloration differences between red-shafted and yellow-shafted flickers. Individuals are scored from 0 (pure yellow-shafted) to 4 (pure red-shafted) for each trait. An overall phenotype score is calculated by summing across the six traits and transforming to range from 0-1 (to allow comparisons between the sexes). Phenotypic scoring is adapted from Short (1965).

| **Phenotype score** | **Description** |
|---|---|
| *Wings and tail ("shaft") color, Carotenoid* | |
| 0 | Bright yellow, as in yellow-shafted |
| 1 | Yellow-orange traces, faint in all feathers or heavy in one or several |
| 2 | Orange to red rachises with yellow-orange vanes |
| 3 | Orange-red |
| 4 | Deep salmon red, as in red-shafted |
| *Nuchal patch presence, Carotenoid* | |
| 0 | Present and broad, as in yellow-shafted |
| 1 | Present and restricted in width (less than one-half of normal width) |
| 2 | Present and broken in one or more places |
| 3 | Traces present, usually at sides of nape |
| 4 | Absent, as in red-shafted |
| *Crown color, Melanin* | |
| 0 | Gray, as in yellow-shafted |
| 1 | Gray with brown traces in forehead and crown |
| 2 | Mixed gray and brown (crown half brown with more gray on hind neck) |
| 3 | Crown brown with hind neck gray toward back |
| 4 | Brown confluent with back color, as in red-shafted |
| *Ear covert color, Melanin* | |
| 0 | Tan, as in yellow-shafted |
| 1 | Tan with gray traces |
| 2 | Mixed gray and tan |
| 3 | Gray with tan traces (especially below eye) |
| 4 | Gray, as in red-shafted |
| *Throat color, Melanin* | |
| 0 | Tan, as in yellow-shafted |
| 1 | Tan with gray traces (usually on lower throat) |
| 2 | Mixed gray and tan |
| 3 | Gray with tan traces (usually near chin) |
| 4 | Gray, as in red-shafted |
| *Malar stripe color (males only), Melanin and Carotenoid* | |
| 0 | Black, as in yellow-shafted |
| 1 | Black with <20% red |
| 2 | Mixed black and red |
| 3 | Red with <20% black |
| 4 | Red, as in red-shafted |

### Supplementary Table 2 | Sample information

| Individual ID | Taxa | Sex | Year | County, State | Phenotype score |
|---|---|---|---|---|---|
| LSU B48981 | Yellow-shafted | Male | 2002 | Brevard, FL | - |
| LSU B48980 | Yellow-shafted | Male | 2002 | Escambia, FL | - |
| LSU B50722 | Yellow-shafted | Male | 2003 | Escambia, FL | - |
| LSU B59061 | Yellow-shafted | Male | 2004 | Escambia, FL | - |
| LSU B59422 | Yellow-shafted | Male | 2005 | Escambia, FL | - |
| CUMV 51231 | Yellow-shafted | Male | 2004 | Tompkins, NY | - |
| CUMV 52455 | Yellow-shafted | Male | 2006 | Tompkins, NY | - |
| CUMV 52999 | Yellow-shafted | Male | 2009 | Tompkins, NY | - |
| CUMV 54562 | Yellow-shafted | Male | 2011 | Tompkins, NY | - |
| CUMV 58977 | Yellow-shafted | Male | 2017 | Tompkins, NY | - |
| 1803-25407 | Hybrid | Male | 2016 | Lincoln, NE | 0.000 |
| CUMV 56730 | Hybrid | Female | 2016 | Butler, NE | 0.050 |
| CUMV 57686 | Hybrid | Male | 2017 | Deuel, NE | 0.083 |
| 1803-25405 | Hybrid | Male | 2016 | Keith, NE | 0.083 |
| 1803-25403 | Hybrid | Male | 2016 | Keith, NE | 0.083 |
| 1803-25410 | Hybrid | Female | 2016 | Sedgwick, CO | 0.100 |
| CUMV 56731 | Hybrid | Male | 2016 | Buffalo, NE | 0.125 |
| CUMV 58091 | Hybrid | Male | 2018 | Keith, NE | 0.125 |
| CUMV 56715 | Hybrid | Female | 2016 | Lancaster, NE | 0.150 |
| CUMV 56717 | Hybrid | Male | 2016 | Buffalo, NE | 0.167 |
| CUMV 58065 | Hybrid | Male | 2018 | Garden, NE | 0.167 |
| CUMV 56728 | Hybrid | Male | 2016 | Keith, NE | 0.167 |
| 1803-25404 | Hybrid | Male | 2016 | Keith, NE | 0.167 |
| CUMV 57607 | Hybrid | Female | 2017 | Logan, CO | 0.200 |
| CUMV 56716 | Hybrid | Female | 2016 | Polk, NE | 0.200 |
| CUMV 58072 | Hybrid | Male | 2018 | Logan, CO | 0.208 |
| CUMV 56725 | Hybrid | Male | 2016 | Morrill, NE | 0.208 |
| CUMV 58060 | Hybrid | Female | 2018 | Morgan, CO | 0.250 |
| 1803-25406 | Hybrid | Male | 2016 | Lincoln, NE | 0.292 |
| CUMV 56724 | Hybrid | Male | 2016 | Morrill, NE | 0.292 |
| CUMV 58090 | Hybrid | Male | 2018 | Morgan, CO | 0.333 |
| CUMV 58084 | Hybrid | Male | 2018 | Logan, CO | 0.375 |
| CUMV 58067 | Hybrid | Male | 2018 | Weld, CO | 0.375 |
| CUMV 57608 | Hybrid | Female | 2017 | Logan, CO | 0.400 |
| CUMV 56734 | Hybrid | Female | 2016 | Morrill, NE | 0.400 |
| 1833-36504 | Hybrid | Female | 2016 | Sedgwick, CO | 0.400 |
| CUMV 58076 | Hybrid | Female | 2018 | Weld, CO | 0.450 |
| CUMV 57988 | Hybrid | Male | 2017 | Morgan, CO | 0.458 |
| CUMV 58148 | Hybrid | Male | 2018 | Garden, NE | 0.500 |
| CUMV 56726 | Hybrid | Male | 2016 | Scotts Bluff, NE | 0.500 |
| CUMV 57610 | Hybrid | Female | 2017 | Logan, CO | 0.550 |
| CUMV 58085 | Hybrid | Female | 2018 | Logan, CO | 0.550 |
| 1833-36502 | Hybrid | Male | 2016 | Morgan, CO | 0.583 |
| CUMV 58079 | Hybrid | Male | 2018 | Morgan, CO | 0.583 |
| 1803-25408 | Hybrid | Female | 2016 | Morgan, CO | 0.600 |
| 1803-25409 | Hybrid | Male | 2016 | Sedgwick, CO | 0.625 |
| CUMV 58068 | Hybrid | Male | 2018 | Scotts Bluff, NE | 0.667 |
| CUMV 57967 | Hybrid | Male | 2017 | Weld, CO | 0.667 |
| CUMV 58080 | Hybrid | Male | 2018 | Morgan, CO | 0.708 |
| CUMV 58069 | Hybrid | Male | 2018 | Weld, CO | 0.708 |
| CUMV 58070 | Hybrid | Male | 2018 | Weld, CO | 0.750 |
| CUMV 56723 | Hybrid | Female | 2016 | Kimball, NE | 0.800 |

| | | | | | |
|---|---|---|---|---|---|
| CUMV 56736 | Hybrid | Male | 2016 | Scotts Bluff, NE | 0.833 |
| CUMV 58078 | Hybrid | Male | 2018 | Larimer, CO | 0.875 |
| CUMV 56727 | Hybrid | Male | 2016 | Scotts Bluff, NE | 0.917 |
| CUMV 58063 | Hybrid | Male | 2018 | Larimer, CO | 0.958 |
| CUMV 58077 | Hybrid | Male | 2018 | Larimer, CO | 0.958 |
| 1833-36503 | Hybrid | Male | 2016 | Larimer, CO | 1.000 |
| BURKE 109367 | Red-shafted | Male | 2002 | Josephine, OR | - |
| BURKE 113386 | Red-shafted | Male | 2002 | Josephine, OR | - |
| BURKE 112778 | Red-shafted | Male | 2002 | Josephine, OR | - |
| BURKE 101882 | Red-shafted | Male | 2002 | Josephine, OR | - |
| BURKE 101883 | Red-shafted | Male | 2002 | Josephine, OR | - |
| BURKE 100969 | Red-shafted | Male | 2003 | Inyo, CA | - |
| LSU B34359 | Red-shafted | Male | 1999 | San Bernardino, CA | - |
| LSU B24273 | Red-shafted | Male | 2000 | San Bernardino, CA | - |
| LSU B60069 | Red-shafted | Male | 2007 | San Bernardino, CA | - |
| BURKE 66173 | Red-shafted | Male | 1996 | Tulare, CA | - |

**Supplementary Table 3 | Genomic regions identified in the GWAs.** List of regions identified as significant in the six genome-wide association (GWA) analyses of hybrid flickers. Each region includes information on the trait (or traits) it was significantly associated with, the number of significant SNPs in the region, and the number of identified genes. Chromosomal and base pair positional information is based on alignment to the zebra finch genome.

**Supplementary Table 4 | Candidate genes.** List of candidate genes within the genomic regions of interest shown in Supplementary Table 3. Gene functions of potential relevance to melanin or carotenoid pigmentation are included (e.g., vesicles, WNT signaling pathway). The strongest candidate genes, those with known or suspected roles in pigmentation, are shaded in gray. Chromosomal and base pair positional information is based on alignment to the zebra finch genome.

**Supplementary Table 5 | Summary statistics averaged across the whole genome**

| Taxa | Observed Heterozygosity | Nucleotide Diversity | Tajima’s D |
|---|---|---|---|
| Red-shafted | 0.3630 | 0.0016 | 0.0843 |
| Yellow-shafted | 0.3979 | 0.0014 | 0.1153 |
| Hybrid | 0.3115 | 0.0015 | 0.6423 |

**Supplementary Fig. 1 | Distribution of red-shafted and yellow-shafted flickers.** Geographical distribution of the red-shafted and yellow-shafted flickers in North America with the approximate location of the hybrid zone shown in orange.

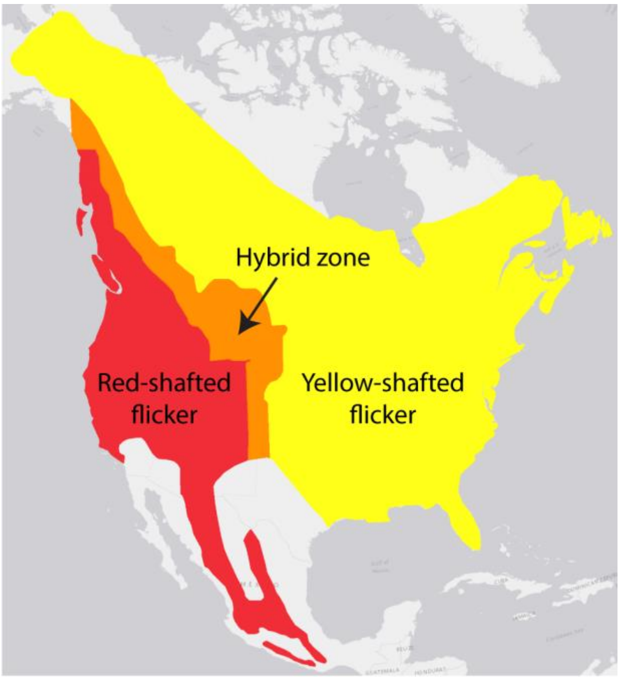

**Supplementary Fig. 2 | Additional PC axes.** PC3 and PC4 of the principal component analysis (PCA) showing the hybrid flickers (orange points) separating from red-shafted (red points) and yellow-shafted (yellow points) flickers on these axes.

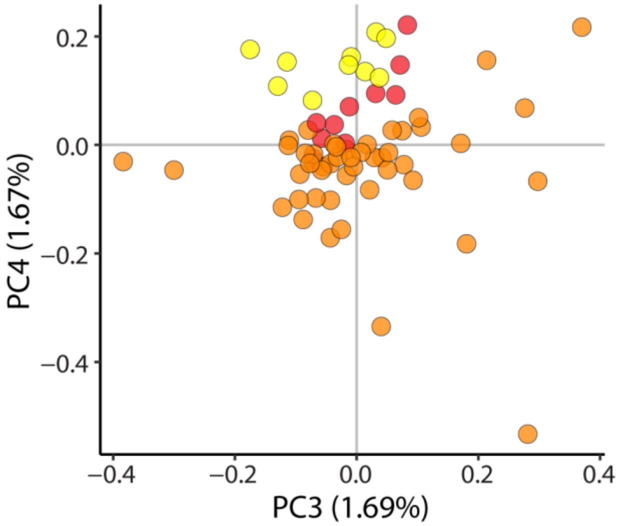

**Supplementary Fig. 3 | PCA of fixed SNPs.** PCA using 790 fixed SNPs ($F_{ST}$ = 1) between allopatric red-shafted and allopatric yellow-shafted flickers with points colored by phenotype score with allopatric samples colored at the ends of the gradient.

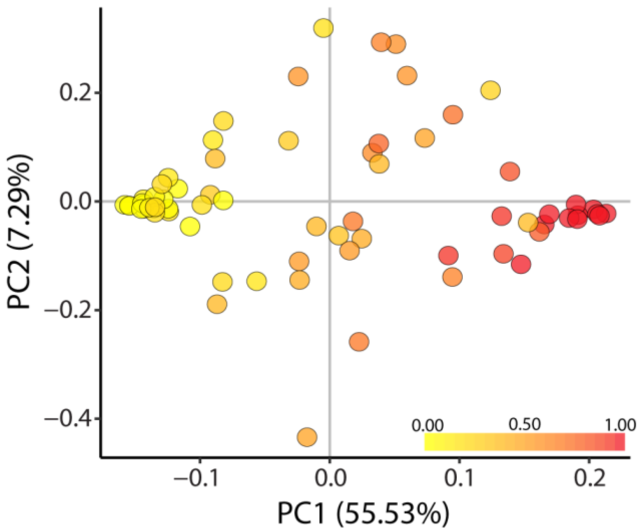

**Supplementary Fig. 4 | Venn diagram of SNPs identified in the GWAs.** Venn diagram showing how individual SNPs identified as significantly associated with coloration traits in the independent GWAs are shared across analyses.

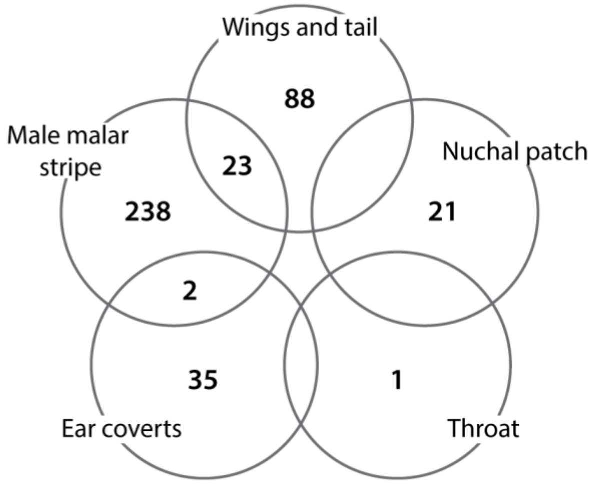

**Supplementary Fig. 5 | Associations between genomic SNPs and randomized phenotypes.** Results from the GWAs comparing individual SNPs with the six plumage patches after the phenotypes were randomized across individuals. For visualization purposes we show only points with $-\log_{10}(P) > 2$.

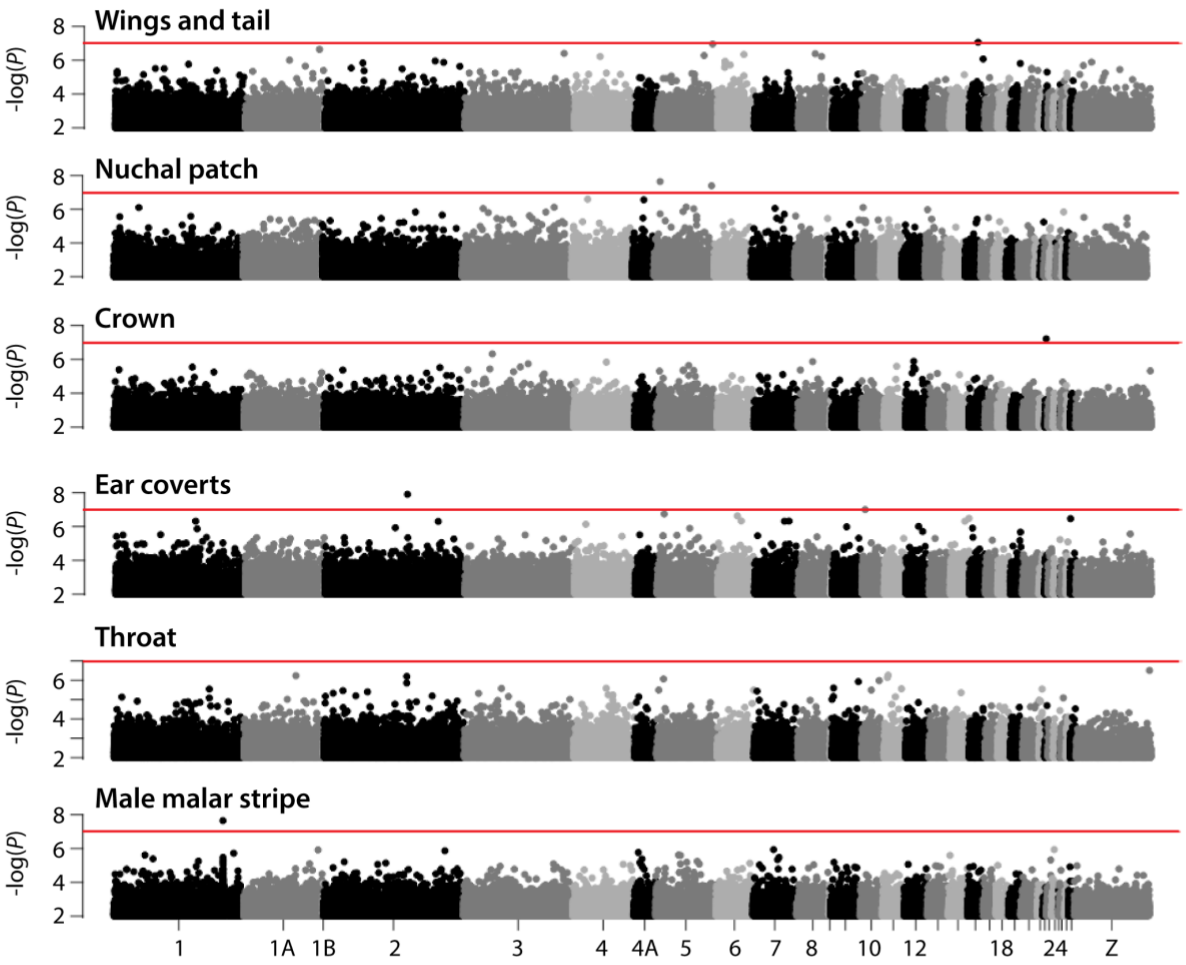

**Supplementary Fig. 6 | *$F_{ST}$* of SNPs identified in the GWAs.** Distribution of per-SNP $F_{ST}$ values of SNPs identified as significantly associated with coloration traits in the GWAs.

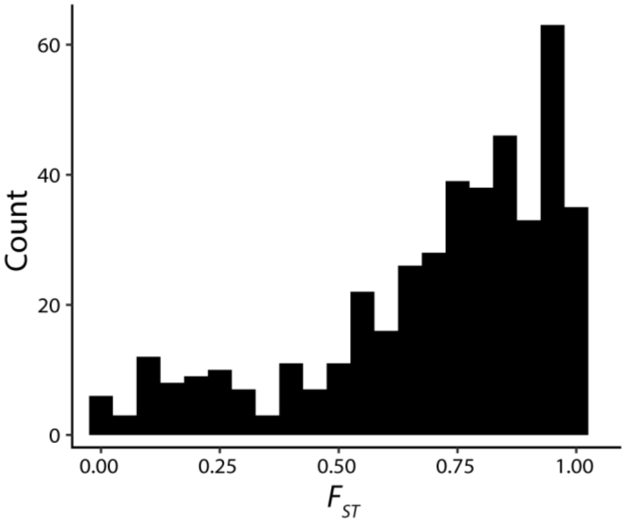
